# The urinary tract microbiome in older women exhibits host genetics and environmental influences

**DOI:** 10.1101/838367

**Authors:** AS Adebayo, G Ackermann, RC Bowyer, P Wells, G Humphreys, R Knight, TD Spector, CJ Steves

## Abstract

The urinary microbiome is a relatively unexplored niche despite the fact that we now know that it is not sterile. Moreover urinary microbes, especially in ageing populations, are associated with morbidity even when infection is subsequently not proven. We present the first large-scale study to explore factors defining urinary microbiome composition in community-dwelling older adult women without clinically active infection. Using 1600 twins, we estimate the contribution of genetic and environmental factors to variation in microbiome using both 16S and shotgun metagenomics. We found that the urinary microbiome is distinct from nearby sites and is unrelated to stool microbiome. Core urinary microbiome taxa were defined. The first component of weighted unifrac was heritable (18%) as were key taxa (e.g *Escherichia-Shigella* (A>0.15)). Age, menopausal status, prior UTI and host genetics were top among factors defining the urobiome. Increased composition was associated with older age, contrary to previous findings.

## Introduction

The resident microbial community (microbiome) at different human body sites, continues to generate research interest, driven by evidence of a role in human physiology. The study of the urinary microbiome (urobiome) is much less established compared to the gut microbiome; perhaps due to the previous belief that the urine was sterile in the absence of a urinary tract infection. Recently, research has shown that this is not the case and that the urinary tract is in fact, another site with a microbiome, reflective of the microbes inhabiting the bladder and closely associated organs (Wolfe et al., 2012; Siddiqui et al., 2012; Whiteside et al.; 2015). This evidence is supported by enhanced quantitative cultures, 16S marker studies and metagenomics, in different populations (e.g Kramer et al., 2018; Adebayo et al., 2017;Wu et al., 2017).

Studies to date have identified differences in the urobiome in relation to urinary tract conditions (Sihra et al., 2018; Wolfe & Brubaker, 2019) including urinary infections (UTI). There is evidence for sex differences in the urinary microbiome which may in part be due to differences in the length of the urinary tract (Moustafa et al. 2018). Women are much more likely to develop UTI, with a lifetime risk of up to 50% (Franco, 2005). UTI is also the commonest reason for antibiotic treatment in adult women, which has implications for urinary and other microbiomes and antimicrobial resistance. Early work has indicated that the non-infected state microbiome may influence resilience to infection. Thus this paper is focused on understanding the major factors defining the urobiome in community dwelling women without active infection.

Recent studies involving urinary/bladder microbiomes have involved relatively small sample sizes (dozens or few hundreds of people) in hospital or clinic attending patients. For instance, results from our literature search (Jan 2015 to September 2018) included case-control studies on elderly/non-elderly patients (Liu et al., 2017; n=100); urinary tract infections (Moustafa et al., 2018; n=112), cancer (Wang et al., 2017; n=65), diabetes, overactive bladder (Wu et al., 2017; Fok et al., 2018,; n=55-126), chronic kidney disease (Kramer et al, 2018; n=41); surgical transplant patients (Rani et al., 2018, n=20); menopause (Curtiss et al., 2018; n=78). Reinforcing this, a recent review (covering studies up to 2016) carried out by Aragon et al. (2018) reported that the sample sizes in urinary microbiome studies varied between 8 to 60 for healthy controls and 10-197 for cases. Their report shows that many studies are commissioned on incontinence, bladder-related and gynaecologic patients. Moreover, many of the urine microbiome studies, either with 16S or shotgun metagenomes, exclude samples with non-detectable/ below detection microbiome. While the assumption maybe that the failure is completely technical, it is unknown if host factors contribute to having ‘extremely-low’ or ‘below detection’ urine microbiome.

Recently, studies of the gut microbiome, have shown a role of host genetics. While Goodrich and colleagues first reported clearly heritable components within the gut microbiome (Goodrich et al.,2014), a finding which a few subsequent studies have also reiterated(Luca et al., 2018), Rothchilds et al reported that environmental factors may largely blur such host genetics factors (Rothchilds et al.,2018). It is unknown if genetic factors are important in the urinary microbiome.

We aimed to characterize the host influence on the urinary tract microbiome in women. Using midstream urine samples from 1600 females in the TwinsUK cohort, this study, perhaps the largest on urinary microbiome so far, reports about the urinary microbiome composition in an average female population of mainly postmenopausal women with no apparent infection. We hypothesized that, in an unselected average population, (1) the inherent core urinary bacterial community could be defined (2) that the urobiome is influenced by host-specific genetic and environmental factors, (3) that some host-specific factors may relate to undetectable microbial biomass in the urine.

## Results

### Urinary microbiome across studies and were distinct from proximal body sites and shared key taxa

Initially, we compared the overall composition of the urinary microbiome to similar datasets from other body sites using the same bioinformatics pipeline, using similar sized datasets of women aged >45(Supplementary Methods & Data1). Alpha diversity in the urine was, on average, reduced relative to the stool and is comparable in two urine and the vaginal datasets (Fig 1A). Stool samples in the majority ordinated separately from urine samples (Fig 1B) (Supplementary Data 1). Repeating these diversity analyses with a separate set of random 100 samples each show similar patterns and significance (SFig1A,B). In paired-sample analysis from TwinsUK (Supplementary Data1), urine microbial taxa separated from stool microbial taxa of the same individual (S1C). There was no clear correlation in the pattern of stool and urine microbiome dissimilarity for the paired samples (either obtained at same time point or not) (Mantel’s r≤0.02, p>0.1) and variance was not homogeneous (Levene paired test p=0.02) (Fig 1C-D, SData1). Thereafter, we examined the TwinsUK urinary microbiome dataset alone.

**Fig 1.**
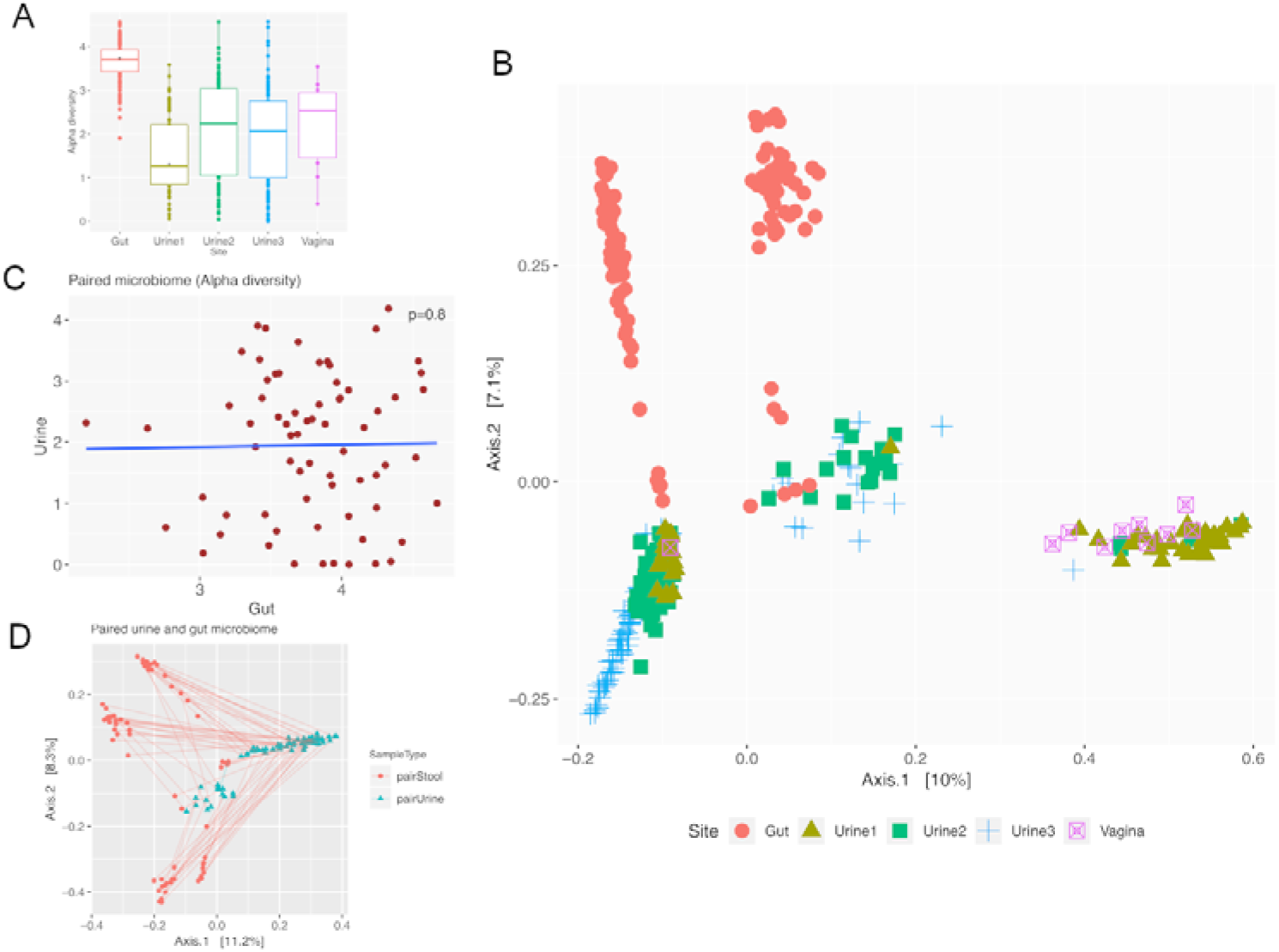
Urinary microbiome in older women is mostly distinct from proximal body sites and unrelated to stool microbiome. **(A) Alpha diversity of urine microbiomes and other body sites.** star symbol indicates significance compared to TwinsUK urobiome. **(B) Dissimilarities in urine microbiomes and other body sites.** Plots are based on unifrac distances **(C) Paired alpha diversity analysis of stool and urine collected at same time point (D) Differences in paired stool and urine microbiome from the same time point.**

### General description of urobiome

Urine samples from 1600 mainly postmenopausal women (mean age 66.4) in the TwinsUK cohort were analysed, revealing 10955 present species-level taxa from filtered 16S data. Participant characteristics are shown in Table 1. There was high level of variability in particular species present in an individual, with only 246 (2.2%) ASVs occurring in at least 5% of samples. The use of a compositionally-sensitive analysis improved the ranking of some abundant taxa as compared to common non-compositional analysis (SFig3). To highlight intra-microbiome relationships, hierarchical balances were created, resulting in mixed-genera subclusters from 61 species-level taxa (hereafter referred to as the core urobiome). There were more Actinobacteria, Fusobacteria and Proteobacteria compared to normal gut microbiome (SFig 3B).

**Table 1.** Summary of participants in TwinsUK urinary microbiome study

| Phenotype category | Subcategory | α-D index (mean±SD) | Ave. no of unique taxa(mean±SD) | No. of samples | Age (mean±SD) |
| --- | --- | --- | --- | --- | --- |
| Participants | | 2.01±1.05 | 65.7±48.8 | 1600 | 66.7±8.3 |
| Previous UTI occurrence<sup>s</sup> | 0 times | 2.14±1.0 | 66.1±43.1 | 393 | 67.6±8.2<sup>s</sup> |
| | 1-4 times | 2.02±1.04 | 67.5±51.0 | 719 | 65.9±7.8 |
| | 5-9 times | 1.98±1.03 | 65.4±45.2 | 208 | 66.3±8.3 |
| | 10times > | 1.79±1.17 | 60.0±53.9 | 201 | 65.7±8.3 |
| Age<sup>s</sup> | <50-54 | 1.56±0.76 | 45.9±32.2 | 117 | - |
| | 55-59 | 1.86±1.13 | 61.7±49.7 | 210 | - |
| | 60-64 | 2.00±0.98 | 63.5±44.8 | 327 | - |
| | 65-69 | 2.04±1.03 | 66.0±49.8 | 409 | - |
| | 70-74 | 2.16±0.97 | 71.5±50.6 | 276 | - |
| | 75-79 | 2.26±1.12 | 74.5±50.1 | 170 | - |
| | 80-84 | 2.02±1.12 | 63.7±41.9 | 68 | - |
| | 85- | 1.73±1.42 | 71.7±62.3 | 23 | - |
| RecentAntibiotic usage:3mths | Yes | 1.97±1.20<sup>ns</sup> | 70.0±53.0<sup>ns</sup> | 47 | 68.3±8.0<sup>ns</sup> |
| | No | 2.03±1.06 | 66.0±49.0 | 945 | 66.6±8.3 |
| Frailty | <0.15 | 2.05±1.01<sup>ns</sup> | 67.0±49.0<sup>ns</sup> | 511 | 65.9±7.5<sup>s</sup> |
| | 0.15-0.29 | 1.99±1.05 | 64.8±49.0 | 834 | 66.1±8.0 |
| | 0.3-0.44 | 2.04±1.15 | 67.5±48.0 | 227 | 68.4±8.9 |
| | >0.45 | 1.75±1.17 | 62.0±47.0 | 28 | 68.5±8.2 |

Legend. $\alpha$ -D: Shannon H index of alpha diversity; No. of taxa refers to number of unique sequence variant per sample i.e. no of potential species. Diversity measures were calculated after subsampling to 2000. S/NS indicates statistical significance or not for tests of a phenotype as a continuous variable. Post-hoc pairwise comparisons showed no difference in alpha diversity after 75years.

Having low reads (no reliably-detected microbiome (<2000 reads post-filtering)) (Supplementary Data 2) associated with younger age and lower level of health deficit; specifically, a ~20% increase in the chances of detectable microbiome for a unit increase in age (p=0.0048, OR=1.21, CI=1.07 - 1.39) and ~14% increase for a unit increase in the frailty index (OR=1.144,CI=1.01-1.30,p=0.0359). There was no association between low read status and the number of previous Urinary Tract Infections (UTIs), recent antibiotics usage, surgery episodes or number of childbirth episodes (parity); amplicon concentrations associated with parity (β=1.89,p=0.0035) alone among other demographics (Supplementary Data 2).

### Host genetics’ influences variation of urine microbiome

First, the quantitative twin model analysis showed considerable and significant genetic component in the first principal coordinate (PCo) of beta diversity (inter-individual) distances which capturing 57% of the variation. Heritability of this first PCo was 18% (A= 0.179, CI=0.05-0.415, p=0.003351; C=0.0049, E=0.8164, n=760 pairs) (Fig 2A). Significant heritability was maintained when adjusting for other factors (Supplementary Data 3). Likewise, treating the microbiome data as Atchinson composition, the first principal component (63% of variation) on inter-sample distances showed 21% heritability (CI=0.10-0.32,C=0.00,E=0.79), and the first PC was also associated with genetic relatedness (family identity) (Kruskal-Wallis p=0.043). Some clusters showed higher heritability (Fig 2B).

**Fig 2.**
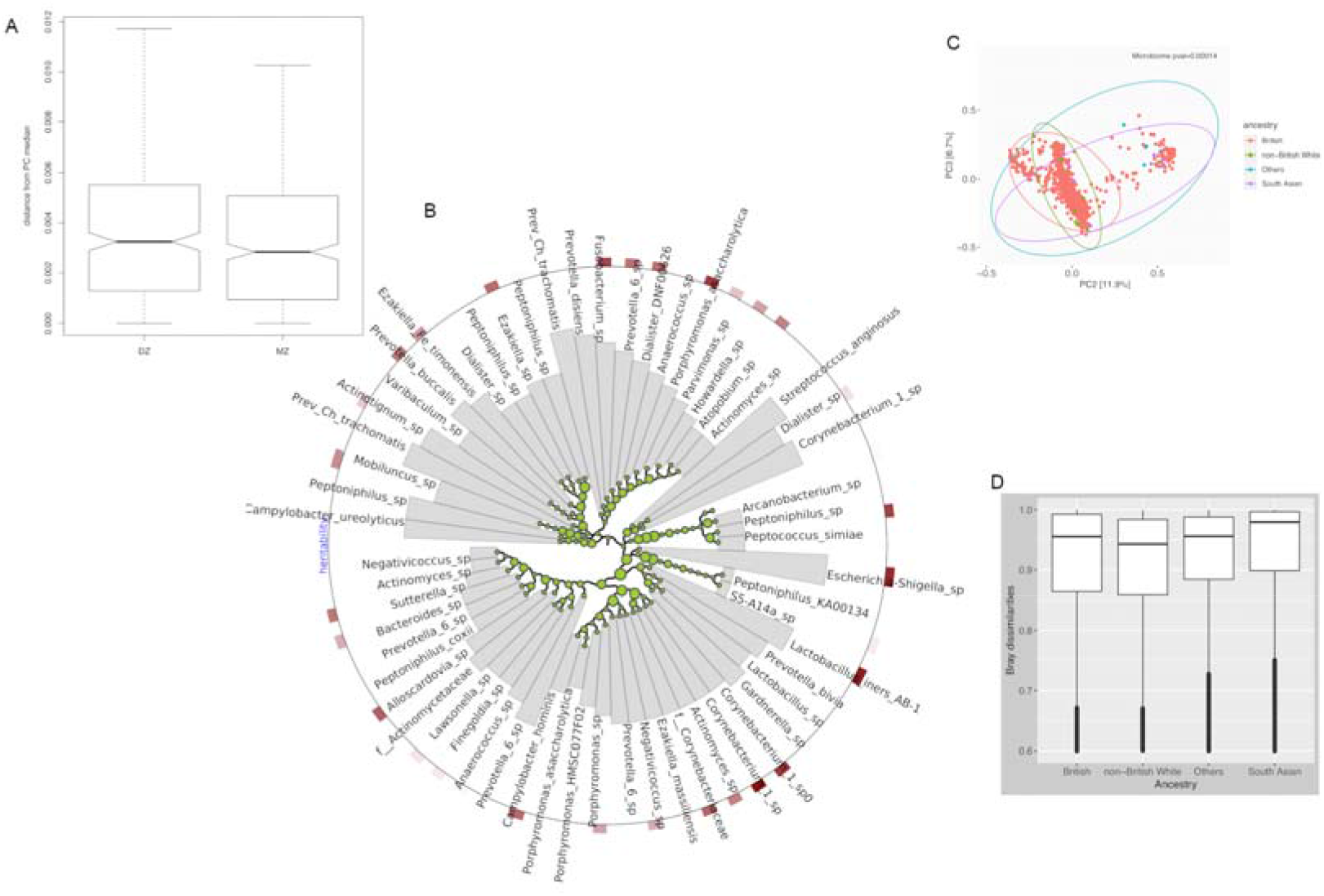
Host genetics considerably influences variation of urine microbiome. **(A) Discordance in paired twin types for Euclidean distances to median microbiome in PC.** MZ-monozygotic pair; DZ:Dizygotic pair; PC: principal coordinate **(B). Heritability and interaction of core urinary microbes.** Size of circles at each subcluster and intensity of rectangular bars at the tips represent increasing heritability of taxa. Neighbouring species in a clade show co-abundance. Taxa are annotated to indicate different species. Clusters are not phylogenetic. **(D). Microbiome principal coordinates with ancestral origin.** White British constitute>90% of individuals. P-values are derived from permutational models due to imbalanced sizes. Ellipses represent 95% confidence interval. **(E) Bray dissimilarities with the ethnic or ancestry divisions.**

In addition, the dissimilarity within relatives (twin pair) in constrained principal coordinates analysis and the average difference in Euclidean distances to the normal PCo median were both smaller for monozygotic pairs (p≤0.027) (Fig 2C and Fig 3D) (Supplementary Data 3), providing further evidence of a genetic component. While the study population was majorly of British ancestry, and therefore ethnicity findings would need to be confirmed, the second PCo of the microbiome diversity differed according to the 4 major ancestry or ethnic origins present (1st PC; p=0.156; 2nd PC p=0.000143), as was the Bray-Curtis dissimilarity between the ancestry groups (Supplementary Data 3, Fig. 2D).

**Fig 3.**
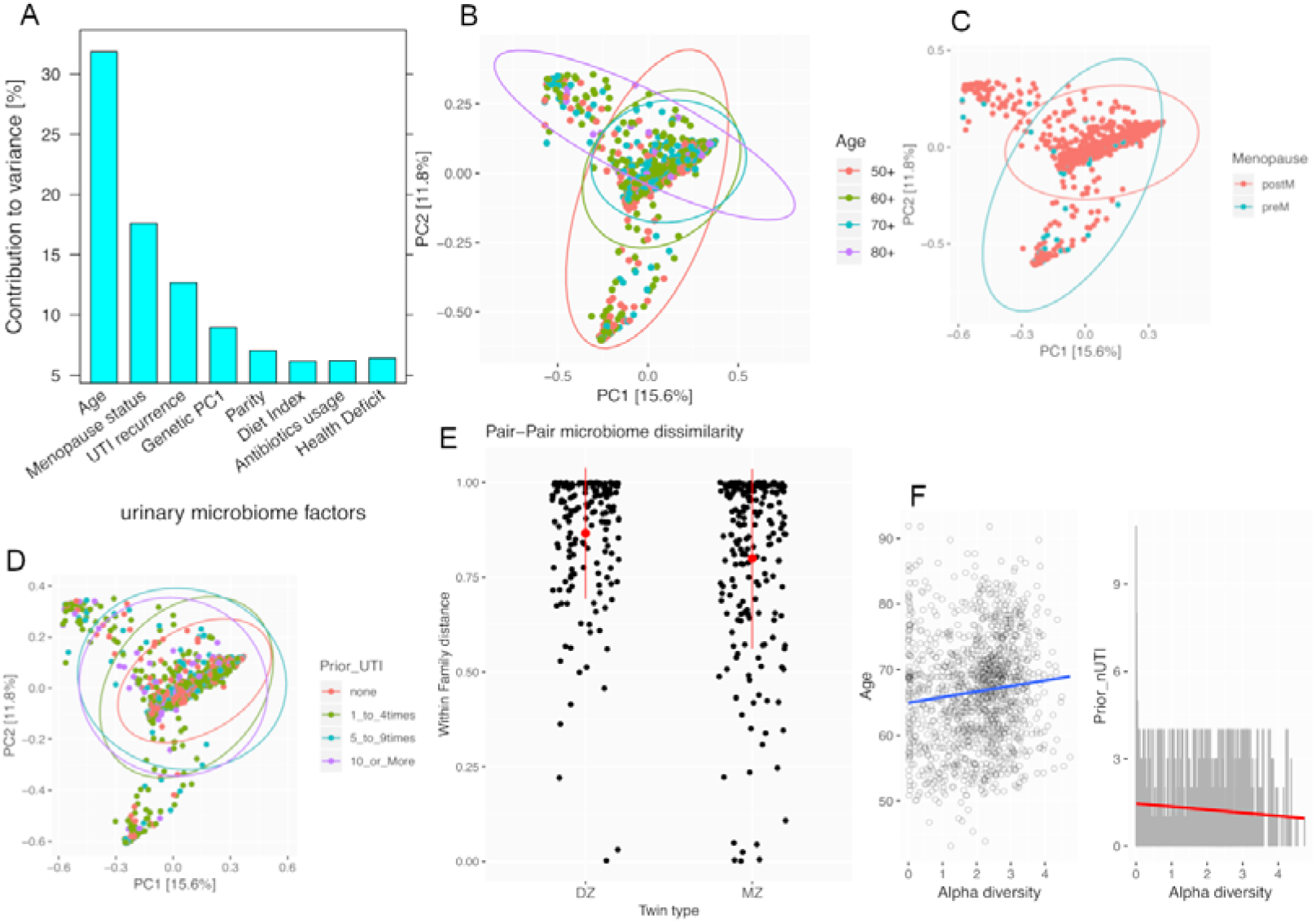
Top contributors to urinary microbiome variation. **(A) Relative contributions to urinary microbiome.** Bars represents average R^2^ for each variable, controlled for the presence of other factors. Microbial variation was measured using Bray-Curtis dissimilarities. Genetic contribution shown was derived from principal components of genetic kinship calculated from whole genome data. **(B-E) Microbiome dissimilarities with (B) age (C) menopause (D) prior number of UTI. (E) within family of twin pairs (F) Trends in intra-individual Shannon diversity with age and prior number of UTI.**

Moreover, the common urobiome taxa (using balance transformations) showed heritability of 23%(95%CI=8.77 to33.7, C=1.66E-12). Almost a quarter (59 of 245) of frequent species had heritability greater than 10%, and some of the most heritable species (e.g. *Lactobacillus iners* AB-1 *and Escherchia-Shigella sp*.) clustered together and members showed phylogenetic relatedness among themselves and with *Christenellaceae* species (SFig 1D-E). Because of the potential role of some of these heritable species in UTIs, we also tested the heritability of occurrence of prior urinary tract infections, finding prior UTI to be significantly heritable (A=0.273, 95%CI=0.178 – 0.368, p=3.073E-13, see Supplementary Data 3) possibly up to 40%.

### Host-related/environmental factors in urinary microbiome, especially age, have important effects

Age, diet, recent antibiotic usage and overall health deficit were assessed in relation to the urobiome as they are known ‘host-specific’ influencers of gut microbiome variation. Parity (previous number of births) and surgical history (had previous surgery or not) were assessed as host-related “environmental” factors as they could potentially alter structures in or proximal to the urinary tract. Previous history of UTI was also assessed.

With increasing age, there is overall increase in alpha diversity (Table 1), which was robust to uneven sample sizes or exclusion of small number of participants aged <50 (0.10≥β≤0.22, 0.00027≤p≥0.0045). Age differed with beta diversity estimates (p<0.001), and was a main influencer of the 3 ‘enterotypes’ (directions) visible in the PCo plot (Fig 2B). The core urobiome and one-third (22) of the subclusters, attained statistical significance with age (1.92E-30≤FDR≤0.046).

The dietary index (the Healthy Eating Index), and an index of health deficit (the frailty index) and antibiotics usage did not produce significant difference in alpha diversity but borderline associations were found with with changes in beta diversity (diet, p=0.052, n=1004; recent antibiotics usage, p=0.041,n=992; health deficit, p=0.031, n=1139). Parity trended toward an association with alpha diversity reduction (p=0.058,n=1047), and significantly with beta diversity (p=0.026,n=1047); surgical history did not differ with beta diversity or alpha diversity (n=540). Occurrence of UTI differed with alpha diversity (p=0.0027) and beta diversity (p=0.001). Similar results were obtained using unifrac sample distances or controlling for other factors.

The contribution to variance that could be attributed to all factors, including host genetics was then examined (Fig 3). For individuals with virtually all phenotypes (n=545), unique contribution was obtained from R^2^ decomposition on microbiome beta diversity estimates, in permutational models (1000 permutations) controlling for other factors. The average for each factor was used after randomly rearranging all factors 20 times. In other scenario of measuring host genetics (Supplementary Data 3), but with a smaller sample size, the contribution of host genetics ranks higher.

### Metagenomes confirm overall 16S microbiome data variation

Using shotgun metagenome data for a subset of 178 individuals, we also examined how closely the overall patterns of the 16S data are replicated in the metagenome data. The classified metagenome reads were 99.64% Bacteria (Supplementary Data 5) and a greater number of urine metagenomes (total and per individual) were obtained than earlier reported in literature. Sample-sample variation or inter-sample distances in the microbiome data were highly correlated from metagenome and 16S data (for Atchinson compositions with Euclidean distance, Mantel’s r=0.859, p=0.002; and for Bray dissimilarities, Mantel’s r=0.799,p=0.001). Sixteen of the top 20 abundant taxa are also within the top 20 of the metagenome data. The core microbiome found in 16S data was largely recapitulated in the metagenomics analysis; 27 of the 31 genera (87%) forming the core urobiome using 16S data were also replicated in the metagenome data. From this core, the total number of species identifiable approximately doubled (125 vs 61 in total, 94 vs 53 in the replicated genera) most likely to due to better species assignment.

## Discussion

In this study, we utilised new approaches in (urinary) microbiome analysis - using amplicon sequence variants rather than OTUs, creating microbial balances from highly frequent taxa, compositional analysis, and eliminating common batch environment effect in twin-pairs - to explore host factors in an relatively large, unselected community-based study population of women. These approaches strengthen deductions made from factors in urinary microbiome variation, for instance, increased diversity with age contrary to previous studies (e.g. Curtiss et al., 2018; Kramer et al., 2018; Liu et al., 2017;Wang et al., 2017).

### Urine and other body sites

The ordination patterns of the microbiomes support current thinking that the urobiome is a distinct site, similar to the observations that most bladder microbiome (urine obtained directly by catheter) differ from vaginal or stool microbiome (Wolfe & Brubaker, 2019). The more divergent of the urine studies (Urine1 cohort) involved patients with incontinence and collection was wholly catheterized. In a very small minority of individuals where urine microbiome taxa appear closer to stool, this is most likely due to phylogenetic or genome similarity in species (as no such closeness occur with non-phylogenetic measures) rather than common demographics (SFig2). In all, the current study show clear dissimilarities in stool and urine for the average population.

### Host-related factors and host genetics’ contribution in urinary microbiome

Parity (childbirth episodes), previous UTI occurrence, recent antibiotics usage and diet showed changes with urine microbiome diversity. Using heritability analysis, the current study showed a considerable genetic influence in the microbiome of ageing women, reaching 15% in 57% of urine taxa variation. The remainder of contribution was largely due to variance unique to individuals. Some clinically important, “uropathogenic” genera such as *Escherichia* had variants with high heritability estimates, In addition, *Lactobacillus. iners*, a commonly found vaginal microbe which is phylogenetically close to the heritable gut microbe Christenellaceae was found to be heritable in urine.

Previously, Rothschild and colleagues (2018) reported that environmental factors such as sharing household may blur genetic influence in gut microbiome composition, while Goodrich and colleagues (2014) showed host genetics played roles in gut microbiome patterns of twin-pairs. The current study, indicates significant contributions of genetics to the pattern of urine microbial composition; and controlling for cohabitation (participants asked if they live together or close with their sibling) and other known factors in urine microbial variation, did not alter the estimated the significant contributions to the pattern. Other parameters from this study bolster the observation on genetic influence: (1) samples of a member in a twin-pair were not extracted or sequenced in the same batch as the other member,(2) adding genetic relatedness statistically explained much more in the pattern of constrained ordination, (3) there was lower intra-twin difference distance to centroid among monozygotic pairs, and (4) there were differences along the lines of ethnic ancestry though the proportion of white British was dominant. Thus we conclude that host genetics influenced variation in urinary microbiome composition in this population of women.

Relative to other factors, only age, menopause status and prior history of current UTI were greater than the influence of genetics. Incidental to our main purpose, we also report here for the first time in humans that history urinary tract infection itself has a significant heritability as suggested in other species (Norris et al., 2000).

### Heritable urinary pathogens

While Corynebacterium species were frequent among top core urobiome taxa with high heritability, the patterns detected for *Lactobacillus iners/jensenii and Escherichia* variants deserve mention. The *Escherichia-Shigella* taxon, renamed as such to reflect the extreme sequence similarity of *Escherichia coli* and *Shigella*, is apparently ubiquitous in the normal urine microbiota from this data. The current study shows that presence of this taxon is influenced by (1) host genetic make up (its proportions had one of the highest heritability estimates (A=0.17,CI=0.11-0.29) of all frequent urine microbial species); and (2) age (its coefficient in age, 0.43, is more than double that of UTI history, 0.20). The relatively high heritability of these taxa were also replicated in the subset with metagenomics data and in all, the findings may have implications in the mixed success of *E. coli* vaccine trials (Huttner et al., 2017).

The current study has limitations. Questionnaire data, which is subject to accurate recall and self-report by participants, was part of measures used in deriving variables such as UTI, diet and frailty. Another limitation may be the use of a single midstream urine sample set from an individual, and as such, prior microbiome stability information is unknown. Clearly, further research is needed to confirm if the findings also relate to the male urinary microbiome.

To conclude, this is the first ‘large-scale’ human study to identify the factors influencing composition of the female urinary microbiome. The urinary microbiome was distinct and apparently unrelated to stool microbiome. It shows a significant contribution of host genetics. Key species known to have pathogenic potential were among the most heritable microbes. Age and menopausal status were the factors with greatest influence on the urinary microbiome in women.

## Acknowledgement

We thank Dr Alan Wolfe and Roberto Limeira of Health Sciences Division, Loyola University, Chicago, United States for providing access to raw sequence data from two urine studies; the phenotype data team at TwinsUK; laboratory team at TwinsUK for sample handling; and Rachel Horsfall, Marina Mora Ortiz, Mary NiLochlainn for discussions and comments on the manuscript. CS received research funding through the Chronic Disease Research Foundation which receives funds from the Denise Coates Foundation. We also thank all participants in TwinsUK (www.twinsuk.ac.uk).

## Author Contributions

Conceptualization: C.J.S, T.S. and A.S.A; Investigation: C.J.S., G.H., G.A., R.B., P.W. and R.K.; Methodology: C.J.S. and A.S.A.; Formal Analysis: C.J.S. and A.S.A; Writing: A.S.A, C.J.S, G.H., T.S. and R.K.; Funding Acquisition: C.J.S. and T.S; Supervision: C.J.S., T.S. and R.K.

## STAR Methods

### 2.1 Cohort and Phenotypes

The TwinsUK cohort has been described elsewhere (Verdi et al. 2019). Participants in the cohort are community dwelling twin pairs, recruited without any specific clinical phenotype. Various demographics were examined. Medical history questionnaires were used to define age (from birth date), history of urinary tract infections (UTIs), cohabitation, antibiotic usage, previous hysterectomy, previous oophorectomy, caesarian section and menopause status. The frailty index, calculated from clinical, physiological and mental domains (Livshits et al., 2017) was used as a measure of health deficit, and the Healthy Eating Index (Bowyer et al. 2018) based on food frequency questionnaires used to assess diet.

### 2.2 16S Microbiome Sequencing and Analysis

Twin-pair samples were separated for processing. Extraction and Sequencing was performed at the Knight Lab, University of California San Diego. A low biomass pipeline designed to extract optimal yields of DNA was used with 16S V4 marker-based paired-end sequencing on IlluminaMiSeq platform. Multilevel quality filtering procedures and data analysis were applied to remove potential contaminants (Suppl Methods). In summary, amplicon sequence variants (ASVs), were filtered, and analysed as individual taxa and as clusters based on highly frequent variants, with subsequent compositional balance transformations (Morton et al.,2017) (Supplementary Methods). The current data was also compared to those of previous microbiome studies with similar age-range of participants after re-analysis of such data to produce ASVs (Supplementary Methods). Diversity analysis was carried out with Shannon index, Unifrac, Bray and Atchinson distances, and permutational analysis of variance was used to test inter-sample differences. Taxa counts were centred-log ratio transformed after adding a pseudocount of 1, and independent taxa associations were pruned for presence in at least 5% of samples.

### 2.3 Metagenome Analysis

Shotgun metagenomic sequencing was carried out for 178 of the participants using newer approaches (Hillman et al., 2018), with additional 14 blanks for quality control. This subset of participants were chosen to include equal numbers of dizygotic pairs and monozygotic twin pairs, as well as equal numbers of twin pairs showing discordance and concordance in 16S microbial diversity. After quality control filtering, and mapped human reads removal (based on hg19) one sample was excluded, and the final analysis included 177 samples, comprising 43 pairs of dizygotic twins and 45 pairs of monozygotic twins (n=176). Potential contaminant species were also removed (Supplementary Methods).

### 2.4 Host genetics analyses

Heritability was calculated using an ACE model in which the component of phenotypes explained by genetics in twin pairs was estimated. Samples from co-twin were separated into different batches for sample preparation and sequencing to remove the shared technical environment related to batching. This further solidified the deductions made from the analysis of the genetic effects. Where constrained principal coordinates analysis was used, microbiome data was ordinated with the family ID tested as a predictor, then the dissimilarity within a family was then extracted to compare twin types. Discordance analysis was based on quantitative difference in pairs of monozygotic and dizygotic twins. Analysis on ethnic origin of participants based on information obtained from questionnaires. To represent host genetic variation, first principal component from SNP-based kinship data, raw whole genomic sequence data (available for a separate subset of unrelated participants) which were part of a previous study (Long et al. 2017), and zygosity:family nested model variance (only for twin-pairs) were obtained. Each of these were analysed separately as a measure of genetic relatedness.

Throughout analysis, technical covariates, including extraction kit lots, mastermix kit lot, batch, extraction and sequencing processors, and depth/library sizes (sequence reads post-QC filtering) were controlled for. Raw sequence data is available from qiita, phenotype data is available on request TwinsUK data access committee at http://twinsuk.ac.uk/resources-for-re-searchers/access-our-data.html. Scripts and codes used are available at github.com/urobiome-host-genetics

## Supplemental Information

**SFig1A-B. Replicate diversity analysis to compare urinary microbiome from various body sites. (C). Plots showing the ordination of paired stool and urine samples (D) Top heritable species.** Species displayed in line bars have more than 15% heritability and star symbol indicate species detected in at least 20%. **(E). Phylogenetic tree of frequent species in urinary microbiome of older women and their heritability.** Tree edges and branch length are coloured by increasing heritability estimates (from green to red). Species displayed in tree were detected at least 5% of study population.

**SFig2. Comparison of demographics for individuals with closer urine and gut microbiome**

**SFig3. Comparison of top abundant urinary microbiome taxa using various approaches**

**SFig4 Additional variation explained from relatedness in twin pairs. A without relatedness B. with relatedness**

