## Supplementary Data and Methods for "The urinary tract microbiome in older women exhibits host genetics and environmental influences"

Supplementary Methods

Filtering and removal of possible contaminant

Initially, common QC processes were followed such as artifact removal, chimera checking, read-length trimming, and short-read discard. As low biomass microbiome tend to be influenced more by contaminants and cross-talk than high biomass sites such as gut, more steps were utilised.

(a)Blank controls (n=105) were sequenced along with normal samples.

(b) Sequence variants (or potential taxa) were removed if the counts attributed to it in the blanks was more than 5% of the total counts for that taxa variant; OR if the number of blanks in which a sequence variant occur is more than 10% of the total number (blanks + actual samples).

(c) Sequence variants were removed if they significantly exhibit a pattern such that its abundance was prevalent in blank controls that were sequenced along with normal samples (e.g. the variant occurs in 50% of blanks but only in 10% of normal samples) or a strong negative correlation (p<0.1) exist between the amplicon library concentration and the number of reads generated for a sequence variant, as implemented in Davis et al. (2018)

Step (b) above was used to complement step c which could not deal with this.

Subsequently, a sample with reads higher than 2000 was deemed to be reliably detected. Setting cut-off at 2000 reads is based on the fact that

(1) It covers about 99.6% of diversity (Shannon) in rarefaction plots and 99.4% coverage (Good’s statistic)

(2) it was much higher than any number reads still present in any blanks after QC and further filtering. i.e. after all QC steps, 30 of 105 blanks sequenced still contained some reads, 90% of these 30 blanks had less than 335 reads, the mean was 152. Because the QC was rather rigorous, these reads are probably due to cross talk in sequencer rather than contaminants.

We also briefly examined potential biological explanations for the occurrence of extremely-low DNA urine sample, apart from efficacy of technical protocols.

Comparison of microbiome studies of similar age

Raw sequence data used in Pearce et al. 2014 (Urine 1), Thomas-White et al. (2017) (Urine 2), Puerto Rico and Plantanal study described as part of Yatsunenko et al. 2012 (Vaginal), Goodrich et al. 2014 (Gut), were obtained on request from authors or from the EBI’s ENA database. These studies, also using 16S V4 region, generally sampled by requesting participants from the general population but some participants in the urine studies were recruited based on specific phenotype of interest. Each sequence set was analysed using the same bioinformatics pipeline described for the current study, and was subsetted to include only women aged 45 years and above to match current study. Re- analysis of these published datasets helped to avoid some data-induced differences in alpha diversity and create a uniform platform for comparison. Also, to minimise multi-study protocol variations and include as many sample as possible, ASV counts were subsampled to 1000 reads in all studies and also subsampled randomly to 100 subjects in each of two replicate sets (except Urine1 and vaginal with smaller participants, n=57 and 11,respectively).

Metagenome Analysis

Paired metagenomics read quality was filtered for average quality value of 30 and merged. Eleven species were removed for presence in blanks and constituting >2% (between 6% and 100%) of the abundance of that species. These ‘contaminant’ species included *Mycobacterium_iranicum, Gordonia_paraffinivorans, Staphylococcus_saprophyticus, Delftia_acidovorans,Corynebacterium_matruchotii,Staphylococcus_capitis, Acinetobacter_harbinensis, Corynebacterium_singulare, Cutibacterium_granulosum, Acinetobacter_towneri, and Cutibacterium_acnes.*

Supplementary Data

Supplementary Data 1

Stool samples had higher alpha diversity than other body sites, (Wilcoxon test, 1.4E-05≤p≥2.2E-16); and urine samples were not statistically different from the vaginal dataset (0.39≤p≥0.72) in alpha diversity except the PM urine study. Differences in the taxa present between individuals were estimated using unifrac beta diversity scores, which takes into account the relatedness of one taxa to another, and these were ordinated in principal coordinates analysis adjusted for study types. Large majority of urine samples shared several related taxa; but a minority were much closer to vaginal (especially from PR urine study). While these described sample separation were for the first two axes and ~20% of variation, the next pair of axes also predict separation patterns on sample type (Fig1C). Repeating the diversity analysis with a separate set of random 100 samples each show similar patterns and significance (S1A,B).

For higher-level changes in classified taxa, each dataset was subsetted to contain the dominant 10 phyla, and their relative abundances across all dataset examined (Fig 1D). The current dataset appear to have more Actinobacteria and Fusobacteria than the other urine datasets (Fig 1D). For paired stool/urine sample analysis, 712 were paired samples from same individual, 69 of which were at same time point. For those at same time point, Mantel r for bray distances (r: -0.02371, p=0.706), weighted unifrac distances (r: 0.1099, p=0.154) and unifrac (Mantel statistic r: -0.06675, p= 0.922).

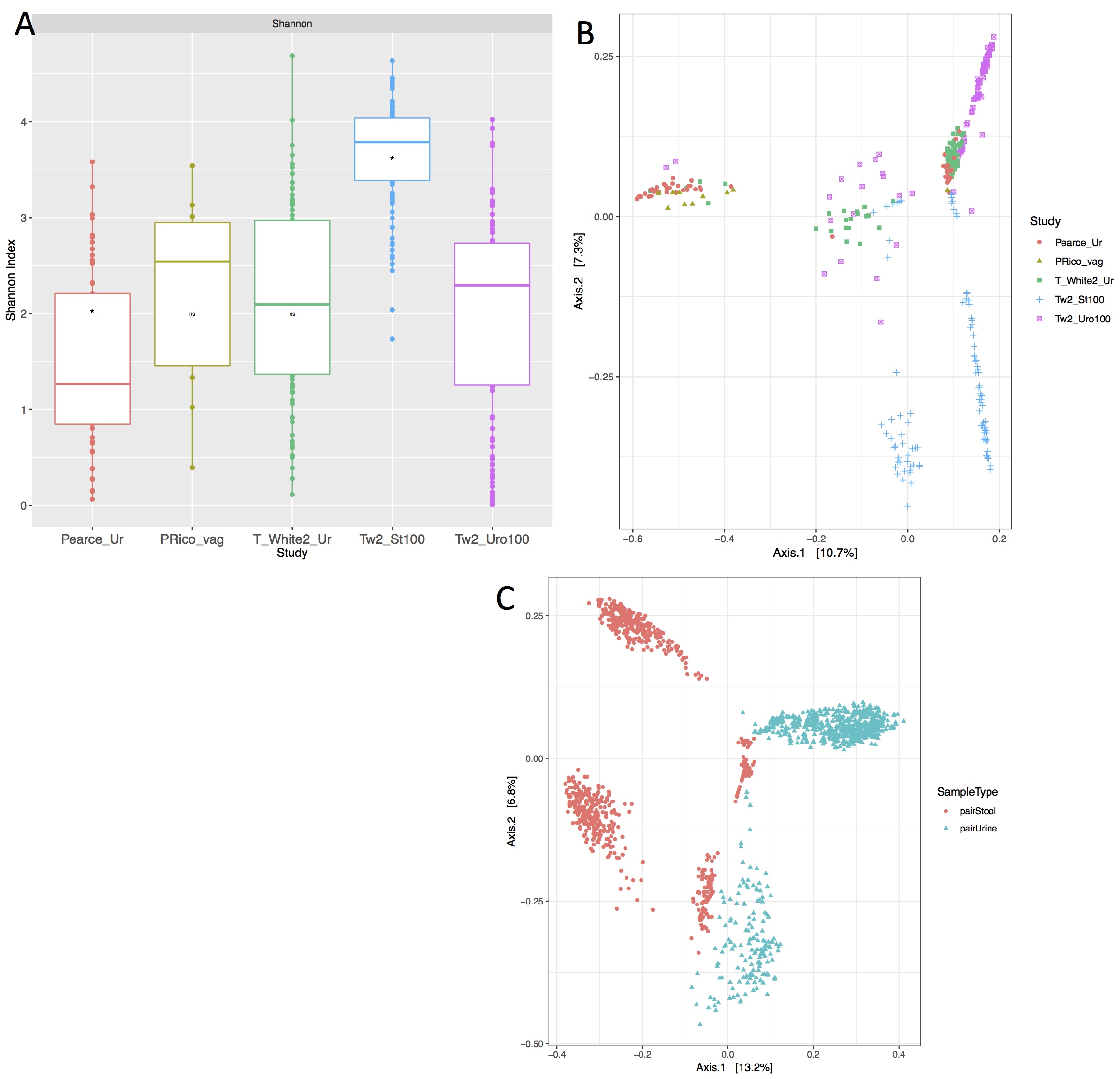

**SFig1A-B. Replicate diversity analysis to compare urinary microbiome from various body sites. (C). Plots showing the ordination of paired stool and urine samples.**

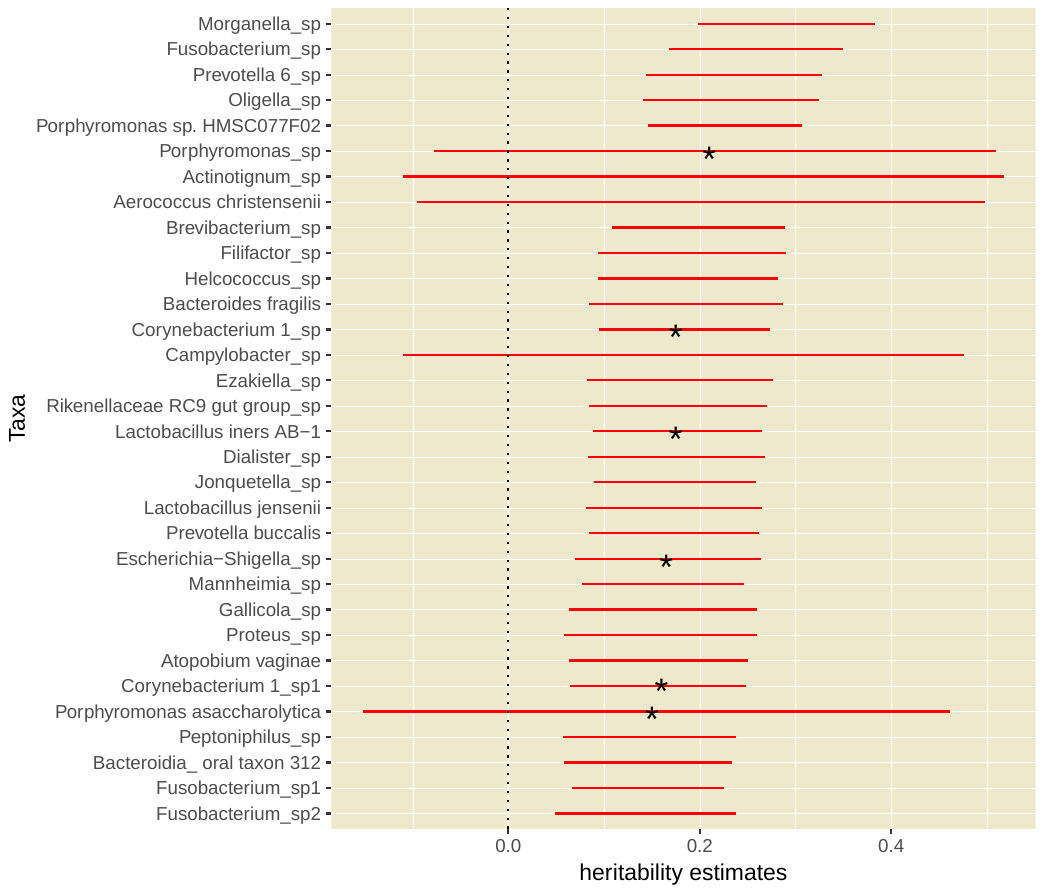

**SFig1 (D) Top heritable species**. Species displayed in line bars have more than 15% heritability and star symbol indicate species detected in at least 20%.

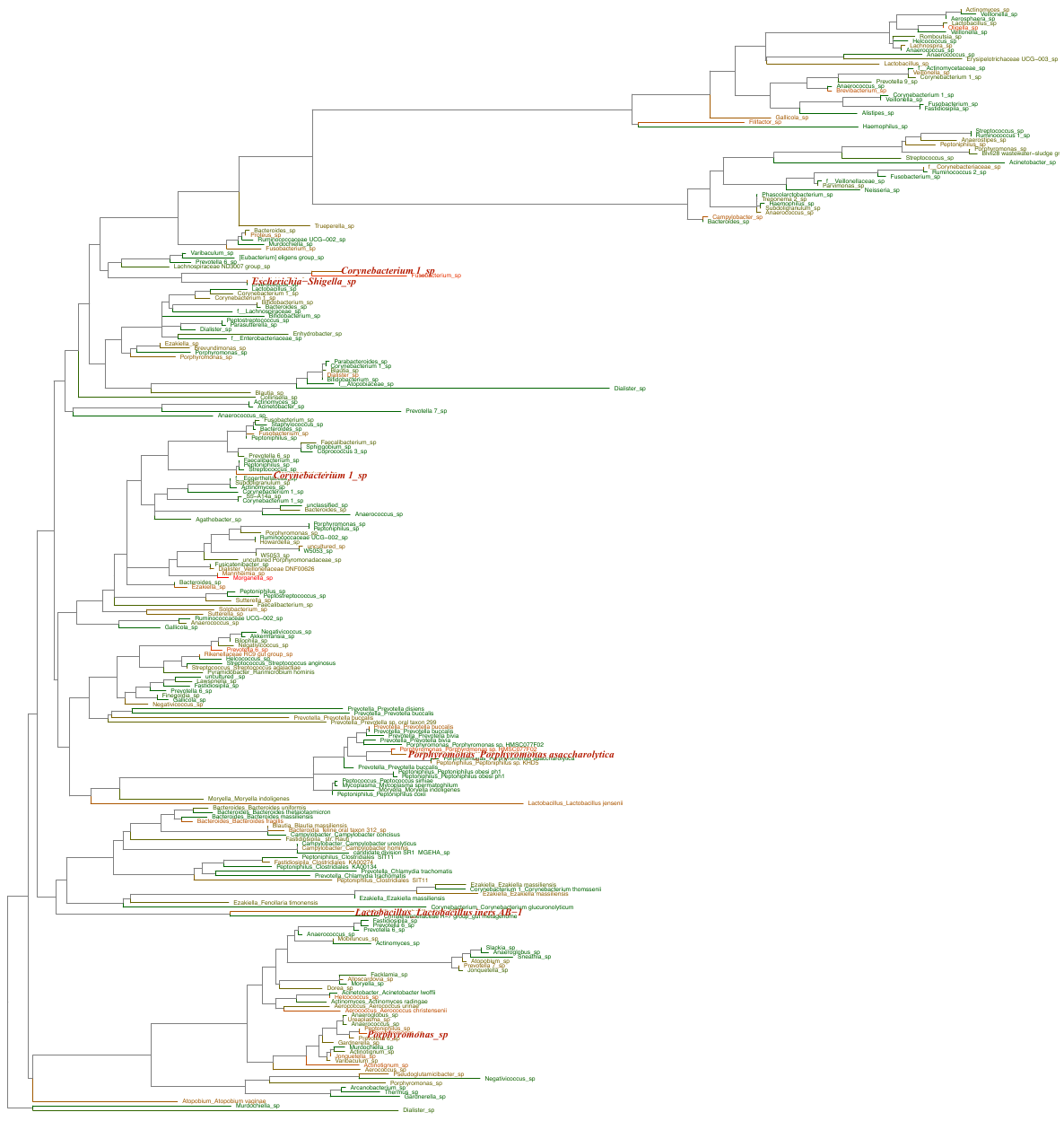

**SFig1 (E). Phylogenetic tree of frequent species in urinary microbiome of older women and their heritability.** Tree edges and branch length are coloured by increasing heritability estimates (from green to red). Species displayed in tree were detected at least 5% of study population.

A minority of urine samples (from various studies) appear to form a different cluster with few stool samples (Fig 1A), hence, we examined summary statistics of such urine samples; and given the recent report of El-Zawahry et al. (2019) on similarities due to biopsy, we also examined pertubations such as surgical (either hysterectomy, oophorectomy or caesarean section) and childbirth episodes. However, we relied more on metadata from the current study as such were not available for samples in the published studies. For these separately-clustered samples, none reported recent antibiotics usage, all were postmenopausal, and from their available data (14), other demographics did not show much difference when compared to the proportions in the rest of the population (SFig2), though slight differences in surgical episodes or births.

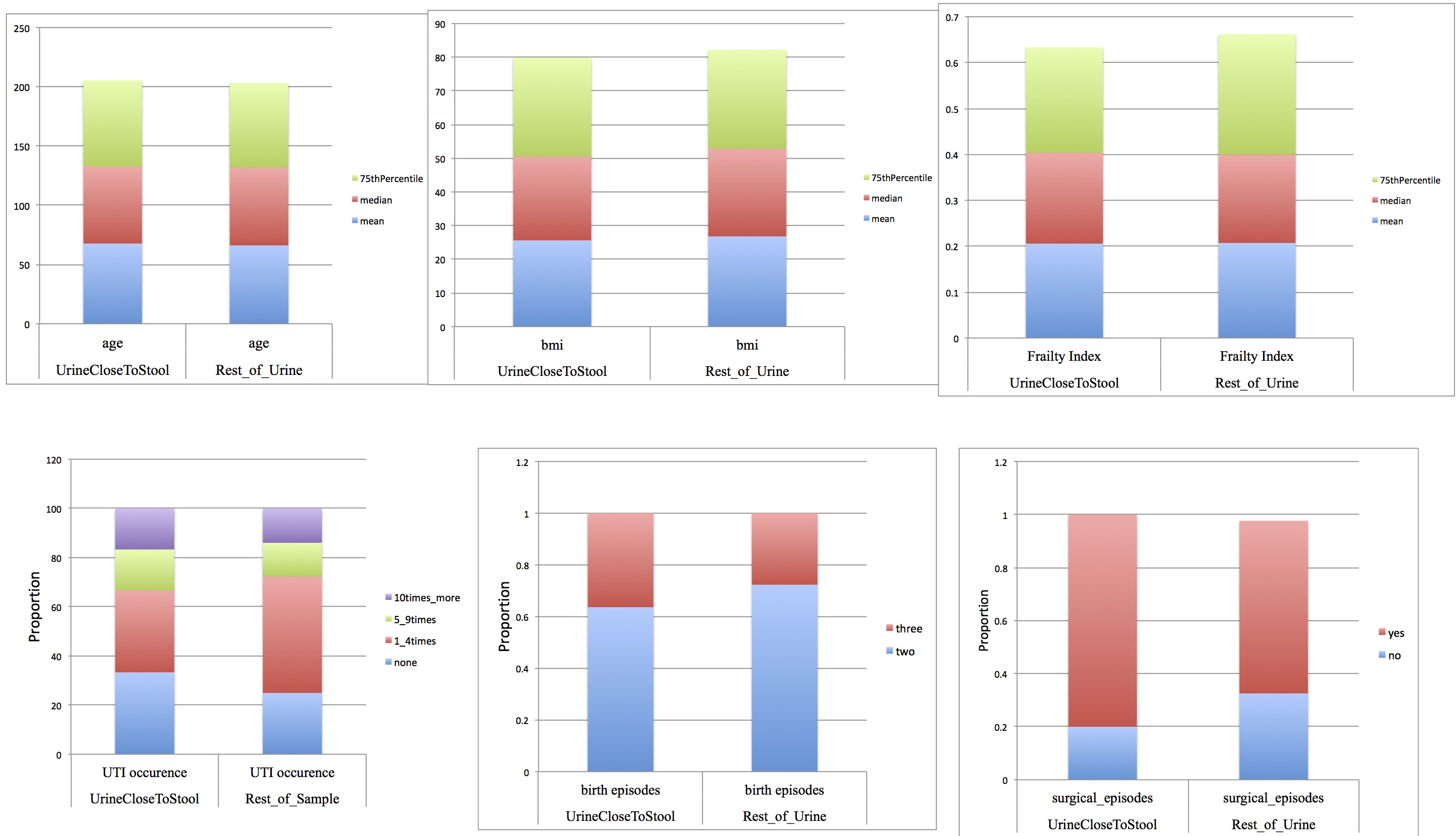

**SFig2. Comparison of demographics for individuals with closer urine and gut microbiome**

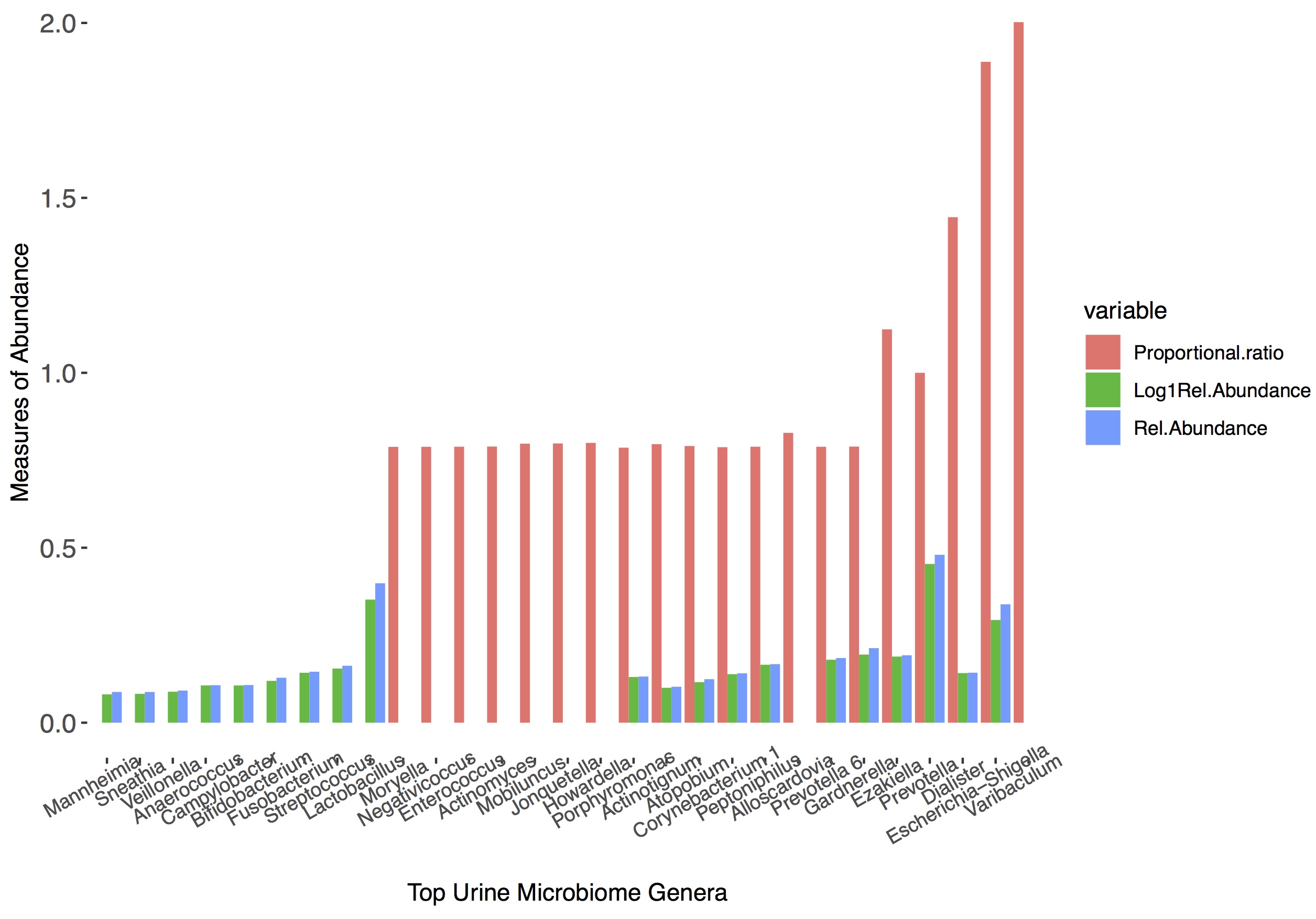

**SFig3. Comparison of top abundant urinary microbiome taxa using various approaches**

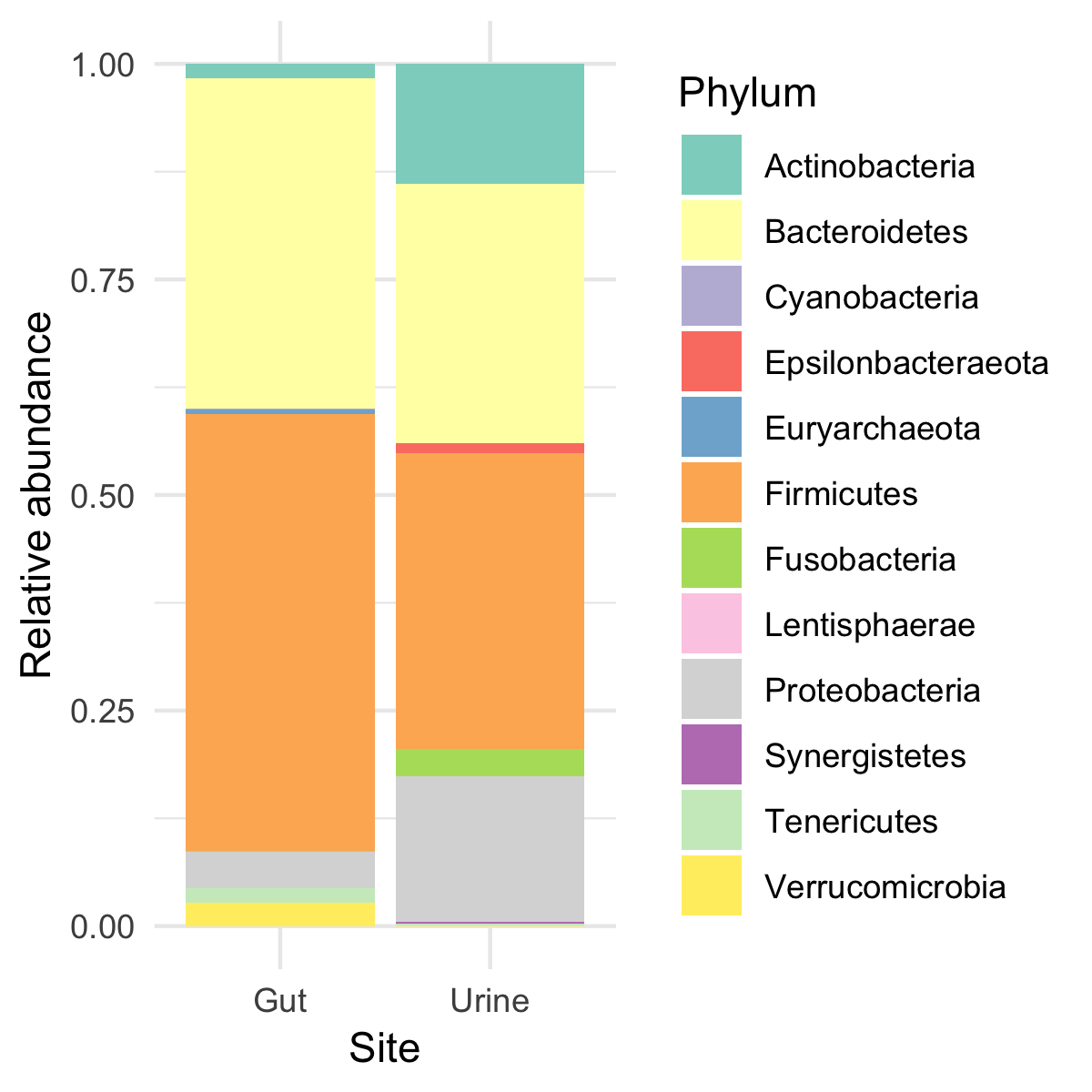

**SFig3B Top microbiome phyla in average population at different body sites.** Gut microbiome data is from 3345 participants and urine microbiome data is from 1230 participants.

Supplementary Data 2

Possible ‘non-technical’ explanations that could affect microbial DNA obtained from urine samples were explored. Controlling for other factors, amplicon concentration associated with parity (number of childbirth) (p=0.0035,β=1.92, n=1346), but not any other variable including age, UTI recurrence, menopause status, health deficit, antibiotics usage, or surgical episodes. Given the highly uneven sample sizes for more than 4 births, permutational tests were also carried out which gave similar results.

Age was slightly lower in those with low-reads (mean age=65.4 ±1SD=7.2, n=370) than those with more than 2000 reads (mean age= 66.7 ±1SD= 8.3, n=1230) (t-test p=0.006). Adjusting for covariates, the odds of obtaining high-read urine samples increased by a factor of 1.19 for an age increase of 1 year (n=1520, p= 0.005,95% CI= 1.06 - 1.35, log odds =0.181). The proportion of those who had no UTI history among the low-read population was 3% more than that of ‘high-read’, the proportions of UTI groups did not differ significantly (χ2 =3.72,p=0.293,n=1521). Although comparing only individuals with no UTI history with ten or more past occurrence, the odds of having a high- read urine sample would increase by a factor of 1.58 (n=594, p=0.0241, 95%CI =1.08 - 2.63, log odds=0.511). There was no heritability in a binary low-read: high-read classification (A=0.0121, 95%CI 0.0086 – 0.156), and discordance was not significant.

Supplementary Data 3

Twin pairs discordance and heritability in microbiome variation

Using principal coordinates analysis on weighted unifrac inter-sample estimates (with species relatedness accounted for) with rarefied data, variance was examined in twin pairs. With all technical covariates controlled for, the unexplained variance among these individuals from the unconstrained residuals were then analysed in 360 twin pairs (207 MZ/173 DZ pairs) in the ACE model. Controlling for age, menopause status, UTI history, and cohabitation, considerable heritability in the singular principal component was (A=0.149, 95% CI: 0.021-0.275, model p 7.88E-6, C=0.0, E=0.8521, n=656, 328 pairs/ 184 MZ/144 DZ). Replicating the analyses based on Bray Curtis metrics rather than the unifrac returned similar estimates at 14.5% heritability (CI=0.03 - 0.269, p=3.561E-06, C=0.0000, E=0.85). The ACE models generally showed substantial unique environmental component but extremely low common environmental component. The next two PCo components (PC2 and 3) capturing 9% and 8% of the variation showed heritability estimates of 6% and 21%.

For the closeness of twin-pairs in ordination, the Euclidean distances to the spatial median sample of the paired twins data was used, with the median based on the initial principal coordinates analysis. The difference for each twin pair was root-normalised distances was smaller for monozygotic pairs than dizygotic pairs (p=0.027, β=-0.005) (Fig2C) and excluding one MZ-pair outlier whose values was more than 10SD.

Also, constrained principal coordinates analysis, which integrates both the calculated diversity estimates and regression, was carried out to deduce the effect of adding relatedness (twin pairs have similar ID) as a factor. It improved the variation explained in just the first 5 axes by almost 12% (first 5= 14.2% compared to 2.4% when constrained only by other covariates) (SFig4).

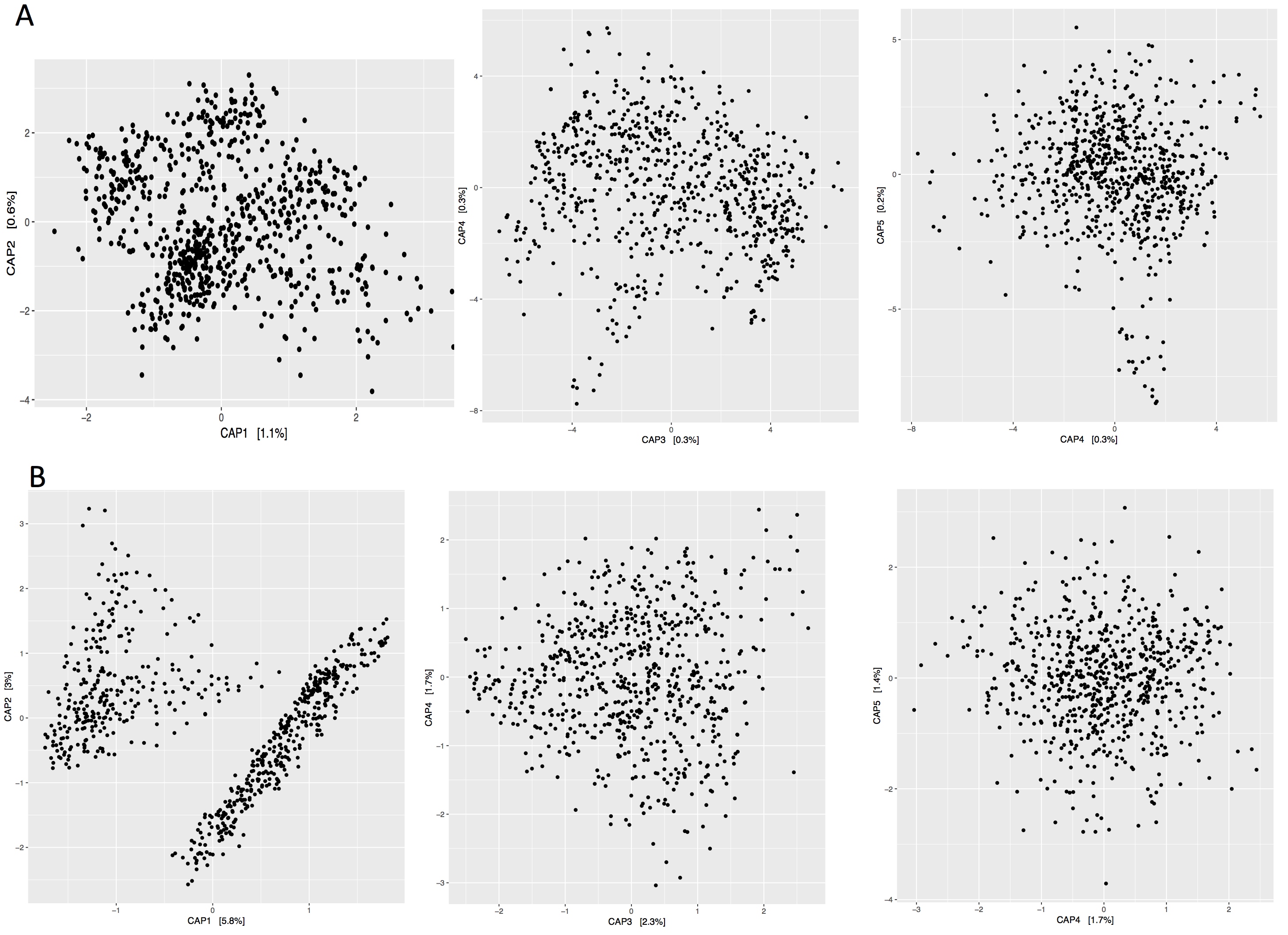

**SFig4 Additional variation explained from relatedness in twin pairs. A without relatedness B. with relatedness**

Ancestry

While the study population had a large percentage of individuals of British ancestry (ethnic English, Welsh, Scottish, Northern Irish & Irish) (n=1391), other ethnic backgrounds: non British white (34), SouthEast Asian (33) and others (1), were among participants. Principal coordinates of the microbiome beta diversity differed according to ancestry (1st PC Kruskal-Wallis; p=0.156; 2nd PC p=0.000143) independent of other factors. To reduce the impact of the sample sizes, permutational anova was carried out, and the microbiome distances was significantly different among ancestry groups (p=0.0026, 9999 permutations) with other factors controlled for.

Heritability in species and in occurrence of UTI

To examine the species or taxa clusters contributing the most to the genetic influence, Normalised ASV abundances in twin pairs were modelled, controlling for covariates, and analysed as clusters and individual species (Fig 2B). The core hierarchical microbial clusters were also analysed using transformed abundances derived from ASVs present in at least 20% of samples were tested and displayed in a tree format (Fig2B).

Having observed (1) that there was genetic influence on the microbiome variation in the current study, (2) the existence of well-known evidence that human genetics play roles in ageing processes and health, and (3) scarce or no literature on the heritability of UTI, we sought to examine genetic influence on the occurrence of urinary tract infections in our study population. The rationale for this was that, as we have presented, the urobiome has heritable features, thus genetic factors could potentially confound the associations. In 573 twin pairs (312 MZ/261DZ pairs, n=1146), UTI history showed heritability (A=0.273, 95%CI=0.178 – 0.368, p<2E-16, C=0.00,E=0.72). Adjusted for cohabitation and age, it gives A=0.265 (CI=0.167-0.359,p=C=0,E=0.73,p<2E-16). Recurrent UTI history (10 or more times) when compared to no UTI history showed a stronger heritability effect (A=0.424,CI=0.248-0.601,p=1.39E-14, C=0.00, E=0.576). This involved 119 twin pairs (67MZ/52DZpairs, n= 238). Adjusted for cohabitation and age retains (A=0.399,CI=0.212-0.586,C=0.00,E=0.601,p=1.38E-12). Additionally, UTI occurrences were scored 0 to 4 from none to 15 times or more, and absolute differences in this score were centered and scaled. Twin-pairs were deemed discordant if a member of the pair had no history and the other had recurrent UTI (10 times or more), and for such pairs, the discordance values were analysed as the ‘distance’ to their twin. Discordance was lower for monozygotic twins than dizygotic (p=0.053,n=486).

Contribution to variance

For twin-pairs with all other factors available (n=310), family of a pair was used as a random effect and as a measure of genetic relatedness of the individuals in linear mixed effect models. Such relatedness was top ranked, explained 22% of variance, and affected the second principal component of microbiome diversity (χ^2^=6.88, p=0.0087). In another scenario, using raw whole genome data from a set of unrelated individuals (Long et al, 2017), some of which were part of the current study, principal components of the genomic variant data were obtained using plink1.9b and the first 10 PC was taken because they each capture just ~1% (total 11%). With all factors inputted (n=178) and average R^2^ taken, in this case, the contribution of host genetics ranked much higher than many other urinary microbiome factors (SFig5)

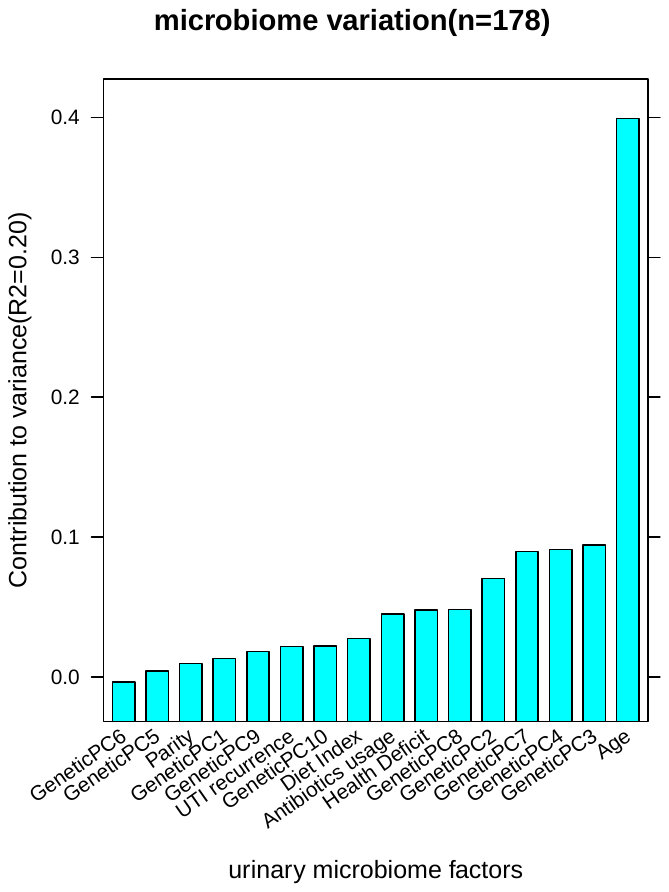

**Contribution of host genetics to urine microbiome variation with host genetic variation taken from top components of genome data of unrelated individuals (n=179)**

Supplementary Data 4

Of the 58.50M reads analysed (excluding bacteria plasmids), Bacteria formed 99.64%, Viruses 0.35%, Archaea 0.0004% in proportion (more bacteria, more viruses and less eukarya compared to Moustafa, 2018). The virome included 224 assigned virus strains, with 19 of them (making up 7.74% of virome abundance) being host viruses (including several polyomavirus, papillomavirus and herpesvirus strains) and others being bacteriophages. Among the human viruses in the urinary virome were the frequent Orgyia pseudotsugata multiple nucleopolyhedrovirus and Glypta fumiferanae ichnovirus, both previously reported only in insect hosts. Median number of metagenomes per individual was 110 species (13-392, mean=121) and 57 genera (11-186,mean=61). Species present in at least 5% of samples were 484, probably high due to full resolution of taxa available for metagenomic sequencing.
